# Seminal quality and global proteomic analysis of spermatozoa from captive Amazon squirrel monkeys (*Saimiri collinsi* Osgood, 1916) during the dry and rainy seasons

**DOI:** 10.1101/771295

**Authors:** Danuza Leite Leão, Sheyla Farhayldes Souza Domingues, Patrícia da Cunha Sousa, Wlaisa Vasconcelos Sampaio, Fábio Roger Vasconcelos, Arlindo Alencar Moura, Regiane Rodrigues dos Santos, Morten Skaugen, Irma Caroline Oskam

## Abstract

The squirrel monkey (*Saimiri collinsi*), a Neotropical primate endemic to the Amazon in Brazil, is used as a biological model for reproductive research on the genus *Saimiri*. Although this animal is known to exhibit reproductive seasonality, nothing is known about the differences in its seminal quality, sperm protein composition, or sperm protein profile between the breeding (dry) and non-breeding (rainy) seasons. Thus, the aims of this study were to evaluate the quality of *S. collinsi* semen during the dry and rainy seasons and to describe the global sperm proteomics and expression variations in the sperm proteins during the two seasons. Aside from the pH, there was no difference in the seminal quality between the dry and rainy seasons. The study approach based on bottom-up proteomics allowed the identification of 2343 proteins present in the sperm samples throughout these two seasons. Of the 79 proteins that were differentially expressed between the two seasons, 39 proteins that were related to spermatogenesis, sperm motility, capacitation, fecundation, and defense systems against oxidative stress were upregulated in the dry season. Knowledge on the sperm proteins provides crucial information for elucidating the underlying mechanisms associated with sperm functionality. Thus, our results help to advance our understanding of the reproductive physiology of *S. collinsi*, providing valuable information for the improvement of protocols used in assisted reproduction techniques for the conservation of endangered *Saimiri* species.

## Introduction

The squirrel monkey (*Saimiri collinsi*), a Neotropical primate endemic to the Amazon in Brazil [1], is commonly used as an experimental model for reproductive research on the genus *Saimiri* [2–4]. According to the International Union for Conservation of Nature’s Red List of Threatened Species, two *Saimiri* species are ranked as vulnerable (*Saimiri oerstedii* and *Saimiri vanzolini*) and one species as almost threatened (*Saimiri ustus*) to extinction [5].

Primates of the genus *Saimiri* exhibit reproductive seasonality. In the free-living animals, the breeding season (mating) and births occur during the dry season and rainy season, respectively. Supposedly, the rainy season is when there is more food available for the newborn [6–8]. However, *Saimiri* monkeys that are held in captivity without variations in their environment and food supply express less of a seasonality pattern by continuing to mate and reproduce throughout the year [9]. Because of the conflicting observations between free-range and captive individuals, it is obvious that the effects of environmental factors (e.g., rainfall, temperature, photoperiod, and food supply) on reproductive seasonality need to be more fully understood [9–11].

Although studies on the squirrel monkey have already shown correlations between reproductive seasonality and spermatogenesis (*Saimiri sciureus*) [12] and the gonadal hormones (*S. sciureus*) [13], only one study has reported the seasonal influence on seminal quality (*S. sciureus*) [14]. However, nothing is known about the protein composition of spermatozoa in these Neotropical primates, or of the differences in the sperm protein profile between the breeding and non-breeding seasons. In domestic animals, proteomic studies have shown the upregulation and downregulation of expression of some sperm proteins when the breeding and non-breeding seasons are compared [15].

Mammalian male fertility depends on physiological events that begin with spermatogenesis and culminate with successful adhesion/signaling between the sperm membrane and the extracellular coat of the oocyte, followed by adhesion/fusion between the oocyte and sperm membranes during fertilization in the female reproductive tract [16, 17]. Proteins expressed by spermatozoa and those from the seminal plasma that bind to the sperm plasma membrane render the spermatozoa capable of fertilizing a mature oocyte [18, 19]. Studies in animals and humans have described sperm proteins that have significant associations with sperm motility (i.e., L-lactate dehydrogenase and dynein heavy chain 1 (DNAH1)) [20, 21], sperm capacitation (i.e., clusterin, spermadhesin, and mitochondrial peroxiredoxin-5) [22, 23], and fertility (i.e., enolase 1, ropporin-1-like protein (ROPN1), and Izumo sperm–egg fusion 1 (IZUMO1)) [24, 25].

In non-human primates, sperm proteomics has been carried out only in Old World primates for characterization of the sperm protein profile [18, 26–29]. Although these studies have been carried out in the genus *Macaca*, which also exhibits reproductive seasonality [30], nothing is known about the changes that may occur in the sperm protein profile during the non-breeding and breeding seasons, and the influence of these changes on the seminal quality of these animals. Knowledge about the absence, presence, underexpression, or overexpression of these sperm proteins could help to further our understanding of the mechanisms behind the reduction in the fertilization ability of sperm [19, 31].

Defining the sperm protein profiles of *Saimiri collinsi* in the breeding (dry season) and non-breeding (rain season) seasons may provide us with a better understanding about the reproductive physiology of these animals, as well as whether the sperm cells could be used in assisted reproduction techniques throughout the year rather than being restricted only to the breeding period. Therefore, the aims of this study were to (i) evaluate the quality of *S. collinsi* semen during the dry and rainy seasons, (ii) describe the global sperm proteomics in *S. collinsi*, (iii) describe the variations of the proteins in sperm collected during the dry and rain seasons, and (iv) evaluate the potential correlation between the expression of the sperm proteins and the seminal quality in *S. collinsi*.

## Methods

### Study design

We conducted a global proteomic analysis of spermatozoa collected from adult squirrel monkeys (*S. collinsi*) throughout an entire year, in the Brazilian Amazon. The seminal coagulum was collected monthly by electroejaculation and liquefied in a powdered coconut water extender (ACP-118; ACP Biotecnologia, Fortaleza, Ceará, Brazil). After 1 h in the ACP-118 extender, the viable sperm cells were separated on Percoll density gradient media and washed. Then, the sperm proteins were extracted and subjected to tryptic digestion, followed by liquid chromatography-tandem mass spectrometry. Statistics and computational biology were used for the identification of the proteins and their relative abundance, categorization of the proteins, and *in silico* analysis of the protein network.

### Animal ethics statement

The animal study was approved by the Ethical Committee in Animal Research (Approval No. 02/2015/CEPAN/IEC/SVS/MS) and by the System of Authorization and Information in Biodiversity (SISBIO/ICMBio/MMA No. 47051-2), and carried the license of the Convention on International Trade in Endangered Species of Wild Fauna and Flora (CITES/IBAMA/Permit No. 17BR025045-DF). All procedures were performed under the supervision of a veterinarian.

### Animals

*S. collinsi* males (N = 4) that originated from the Marajó Archipelago (0°58ʹS and 49°34ʹW) and were maintained in captivity at the Centro Nacional de Primatas, Brazil (1°22ʹ58ʺS and 48°22ʹ51ʺW) were used for the semen collection. The average age of the animals was 15 years. The external genitalia of each animal were evaluated and an andrology examination (i.e., inspection and palpation of the testes to verify the size, consistency, and symmetry) was performed.

### Housing conditions

The animals were housed collectively in cages (4.74 m × 1.45 m × 2.26 m), with 12 h of natural light each day. The mixed animal groups typically consisted of three males and three females and their juvenile offspring. The region is defined by the Köppen-Geiger climate classification system as having a tropical rainforest climate (AF), with an average annual temperature of 28°C (maximum of 32°C and minimum of 24°C) [32]. The animals were fed fresh fruits, vegetables, commercial pellet chow specific for Neotropical non-human primates (18% protein, 6.5% fiber; Megazoo, Minas Gerais, Brazil), and cricket larvae (*Zophobas morio*). Vitamins, minerals, and eggs were supplied once a week, and water was available *ad libitum*.

### Body weight, testicular biometry, and semen collection

Semen was collected monthly from June 2015 to May 2016, every morning before feeding, making up a total of 48 semen collections (12 per animal). For the semen collection, physical restraint of each animal was performed by a trained animal caretaker wearing leather gloves, and all animals were anesthetized with ketamine hydrochloride (20 mg/kg; intramuscularly (IM); Vetanarcol, König S.A., Avellaneda, Argentina) and xylazine hydrochloride (1 mg/kg; IM; Kensol, König S.A.) and monitored by a veterinarian. After anesthesia, the animals were weighed using a weight balance, and the testicular length, width, height, and circumference were measured using a universal caliper. The testicular volume was calculated according to the method described by Oliveira et al. [4]. After the animal had been placed in dorsal recumbency, the genital region was sanitized with a mild soap and distilled water (1:10) and the prepuce was retracted for a more efficient cleaning of the penis with saline solution. The animal was then stimulated according to the rectal electroejaculation procedure described by Oliveira et al. [2–4]. In brief, an electroejaculator (Autojac-Neovet, Uberaba, Brazil) rectal probe was smeared with a sterile lubricant gel (KY Jelly, Johnson & Johnson Co., Arlington, TX, USA) and introduced into the rectum (∼2.5 cm deep) and electrical stimuli were then delivered. The stimulation session consisted of three series (7 and 8 min), composed of 35 electrical stimuli (12.5 and 100 mA), with an interval of 30 s between the series. The ejaculates (liquid and coagulated fractions) were collected into microtubes (1.5 mL).

### Semen evaluation

The 1.5-mL conical microtubes containing the semen were placed in a water bath at 37°C immediately after collection for evaluation of the seminal volume, color, and viscosity. The volumes of the liquid and coagulated fractions were evaluated in a graduated tube, with the aid of a pipette. The appearance was assessed subjectively for color (colorless, yellowish, or whitish) and opacity (opaque or transparent) [2–4]. The seminal pH was measured with a pH strip (Merck Pharmaceuticals, Darmstadt, Germany).

The sperm motility, vigor, and morphology were evaluated according to the methods described by Oliveira et al. [2–4]. For evaluation of the normal sperm morphology and plasma membrane integrity, a smear sample was prepared by adding 5 µL of 1% eosin (Vetec, Rio de Janeiro, Brazil) and 5 µL of 1% nigrosine (Vetec, Rio de Janeiro, Brazil) to 5 µL of semen on a prewarmed (37°C) glass slide. The sperm concentration was determined in a Neubauer chamber after the dilution of 1 µL of semen in 99 µL of 10% formalin solution. The plasma membrane functionality was assessed with the hypoosmotic swelling test after the dilution of 5 µL of semen in 45 µL of hypoosmotic solution (0.73 g of sodium citrate, 1.35 g of fructose, and 100 mL of ultrapure water; pH 7.2 and 108 mOsm/L). After a 45-min incubation in a water bath (37°C), 10 µL of this solution was placed on a prewarmed (37°C) glass slide and covered with a coverslip, and at least 200 spermatozoa were counted to determine the number with coiled tails (indicative of spermatozoa with a functional plasma membrane). All evaluations were performed under a light microscope (E400; Nikon, Tokyo, Japan) at a magnification of 100×. The semen was assessed directly both after collection (fresh) and after dilution in ACP-118.

### Sperm separation and freeze-drying

Owing to the occurance of seminal coagulation in *S. collinsi*, the semen sample was diluted 1:1 in ACP-118 (300 mOsm/kg and pH 6.42), incubated in a water bath (Biomatic, Porto Alegre, Rio Grande do Sul, Brazil) at 37°C for 1 h, and then separated on 45%/90% Percoll gradient media (centrifugation at 10,000 *g*, 15 min, 12°C). Thereafter, the samples were washed in Tris-NaCl medium (centrifugation at 8000 *g*, 5 min, 12°C), and the separated sperm fraction (pellet) was stored in microtubes, together with Tris-NaCl and a protease inhibitor (1:1000; P8340 catalog, Sigma-Aldrich, St. Louis, MO, USA), in liquid nitrogen or a –80°C freezer. For lyophilization, the frozen sperm samples were placed in a freeze dryer (FreeZone 2.5 Liter Benchtop Freeze Dry System; Labconco, Kansas City, MO, USA) for 10 h at a temperature of –55°C and vacuum pressure of 0.025 mbar.

### Liquid chromatography-mass spectrometry

Each individual dried sperm sample was resuspended in 50 µL of lysis buffer (0.1 M Tris-Cl (pH 8.0), 4% sodium dodecyl sulfate, and 10 mM dithiothreitol) and centrifuged at 5000 *g* for 1 h at 4°C. The supernatant was reserved for the preparation of suspension samples for bottom-up proteomic analysis with tryptic digestion, using the method established by Zougman et al. [33]. The extracted peptides were analyzed on an UltiMate 3000 RSLCnano/Q-Exactive system (Thermo Fisher Scientific, Bremen, Germany) that was set up with a Nanospray Flex ion source. The tryptic peptides (∼1 µg loaded) were separated on a 50 cm × 75 µm (i.d.) column (Thermo Fisher Scientific) using a 120 min gradient of 12–45% acetonitrile. The mass spectrometry (MS) and tandem mass spectrometry (MS/MS) data were recorded using a standard data-dependent acquisition method, with the following conditions: *m/z* range of 300–1600; Automatic Gain Control targets of 3 × 10^6^ (MS) and 5 × 10^4^ (MS/MS); resolutions of 70 K (MS) and 35 K (MS/MS); dynamic exclusion set to 20 s, and normalized collision energy set to 28. Xcalibur software (v. 3.1; Thermo Fisher Scientific) was used to evaluate the raw data, which were converted to *mgf* format (for Mascot database searching) using the MS convert module of ProteoWizard (v. 3.0.9016). The Mascot (v. 2.6) searches were performed on an in-house server against an online *Saimiri boliviensis boliviensis* (Bolivian squirrel monkey) database (National Center for Biotechnology Information, Bethesda, MD, USA). MaxQuant software (v. 1.6.1.0) [34] was used for the label-free quantification.

### Protein categorization

The protein information obtained by Mascot was analyzed using the STRuctural Analysis Programs (STRAP) for searching annotations of proteins. STRAP automatically obtains Gene Ontology (GO) terms associated with proteins in an identification list of results based on homology search analysis using various freely accessible databases [335].

### *In silico* protein network analysis

Protein–protein networks were retrieved from the STRING database (v. 10.0), which consists of known and predicted protein interactions collected from direct (physical) and indirect (functional) associations. The database quantitatively integrates interaction data from four sources: a genomic context, high-throughput experiments, co-expression data, and previous knowledge from research publications [36]. The STRING program was set to show no more than 10 interactions and medium confidence. Pathways not described for *S. boliviensis boliviensis* were analyzed using those for other non-human primate species and *Homo sapiens*.

### Statistical analysis

All seminal quality data are expressed as the mean ± standard error of the mean and were analyzed using the StatView 5.0 program (SAS Institute Inc., Cary, NC, USA). Data were checked for normality using the Kolmogorov-Smirnov test. The effects of the dry and rainy seasons on the seminal quality were evaluated by analysis of variance, and differences were determined with Fisher’s protected least significant difference *post hoc* test. A *p* value of <0.05 was considered as being statistically significant. With regard to the differences in protein expression between the dry and rainy seasons, the protein concentration data were logarithmically transformed and two-sample tests were performed using Perseus software (v. 1.6.1.1; Max Planck Institute of Biochemistry, Planegg, Germany).

## Results

### Characteristics of the Amazon monkeys and semen

The body weights of the male monkeys and their total testicular volumes were significantly higher in the rainy season (883.15 ± 14.50 g and 2.42 ± 0.11 cm^3^, respectively) than in the dry season (816.10 ± 6.85 g and 1.91 ± 0.13 cm^3^) (Table 1). Semen collection was successful in 42 of the 48 attempts (88%) because four ejaculates did not contain sperm; of these, 39 samples were used for the experiments. The highest percentage of ejaculates in both the liquid and coagulated fractions was 59%. With regard to the semen color and opacity, 10% of the samples were colorless, 33% were whitish, 57% were yellowish, 46% were transparent, and 54% were opaque. There was a statistical difference (*p* = 0.0002) in the seminal pH between the dry (7.96 ± 0.10) and rainy (7.30 ± 0.11) seasons in the liquid fraction (fresh sample). With regard to the other seminal parameters, there were no changes in the seminal volume, total sperm count, and sperm motility, vigor, plasma membrane functionality, and integrity as well as in the normal sperm regardless of the period of the year (dry or rainy season) (Table 1).

**Table 1.** Mean (±SEM) values of the body weight, testicular volume (cm^3^), and seminal parameters of *Saimiri collinsi* during the dry (breeding) and rainy (non-breeding) seasons.

|  | Dry season | Rain season | P-value |
| --- | --- | --- | --- |
| Body weight (g) | 816.10±6.85 | 883.15±14.50 | 0.0002 |
| **Testicular volume (cm<sup>3</sup>)** |  |  |  |
| Right testicular volume | 0.95±0.06 | 1.19±0.07 | 0.02 |
| Left testicular volume | 0.96±0.07 | 1.22±0.08 | 0.03 |
| Total testicular volume | 1.91±0.13 | 2.42±0.11 | 0.01 |
| **Fresh sample** |  |  |  |
| **Seminal parameters** |  |  |  |
| pH | 7.96±0.10 | 7.30±0.11 | 0.0002 |
| Seminal volume (μL) | 114.70±16.93 | 145.29±28.14 | 0.35 |
| Total sperm count ($\times 10^6$/ml) | 14,196±3,08 | 19,036±8,89 | 0.67 |
| Motility | 63±7.56 | 80±5.39 | 0.76 |
| Vigour | 3.4±0.28 | 3.85±0.28 | 0.23 |
| PMF<sup>1</sup> | 74±2.17 | 73.25±4.67 | 0.87 |
| PMI<sup>2</sup> | 64±5.45 | 59.28±6.94 | 0.59 |
| Normal sperm | 71.40±2.85 | 71.64±5.34 | 0.83 |
| **After dilution in ACP-118** |  |  |  |
| **Seminal parameters** |  |  |  |
| Motility | 44.18±7.82 | 63.46±7.99 | 0.10 |
| Vigour | 3±0.24 | 3.15±0.29 | 0.89 |
| *PMF | 67±4.14 | 73.60±8.41 | 0.46 |
| **PMI | 77.66±3.74 | 67.76±7.62 | 0.26 |
| Normal sperm | 72.58±2.71 | 70.75±5.14 | 0.76 |

\*Plasma membrane functionality (PMF; %)
\*\*Plasma membrane integrity (PMI; %)

### Sperm proteomics

The study approach based on bottom-up proteomics allowed the identification of 2343 proteins in the sperm samples (Supporting Information S1 Table). Of the total proteins identified, 223 were determined to participate in important reproductive events, such as spermatogenesis (67 proteins), sperm motility (42 proteins), capacitation/acrosome reaction (20 proteins), and fertilization (32 proteins) (Supporting Information S2 Table).

On the basis of the GO analysis, the proteins were grouped according to biological process, molecular function, and cellular component (i.e., localization) classes (Fig 1). In the cellular component class, most of the proteins identified were associated with the cytoplasm (12.3%), cytoskeleton (9.4%), and nucleus (8.9%) (Fig 1A). The most common biological processes associated with the proteins were cellular processes (41.6%), regulation (17.6%), and metabolic processes (11.4%) (Fig 1B). Binding (42.8%) and catalytic activity (42.9%) corresponded to the most frequent molecular functions for the proteins (Fig 1C).

**Fig 1.**
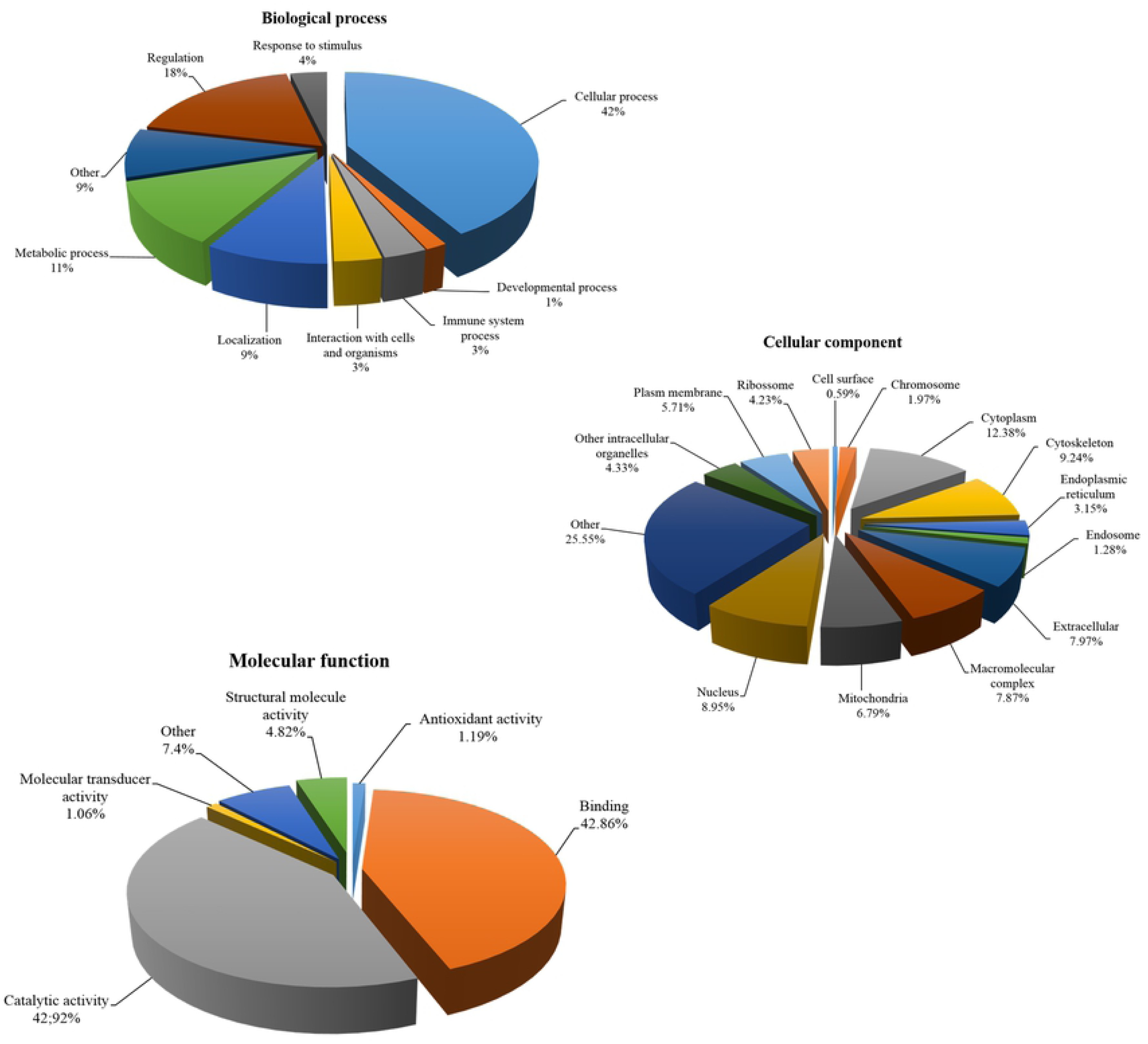
Gene Ontology annotation of the cellular component (A), biological process (B), and molecular function (C) classes of identified *Saimiri collinsi* sperm proteins analyzed by STRAP. The Gene Ontology terms were obtained from the UniProtKB database.

We also identified 79 sperm proteins that were differentially expressed between the dry (breeding season) and rainy seasons (non-breeding season). Of these, 39 were upregulated in the dry season, with the main protein functions being for enzymatic activity (i.e., deoxyguanosine kinase and matrix metalloproteinase-7), cellular regulation (i.e., amine oxidase and serine protease 30-like), and immune system processes (i.e., heat shock 70 kDa protein 1A/1B and clusterin) (Table 2 and Supporting Information S3 Table). With regard to proteins that participate in important events in reproduction, 10 that were increased during the dry season were related to spermatogenesis (i.e., cat eye syndrome critical region protein 5, heat shock-related 70 kDa protein 2 (Hsp70.2/HSPA2), and peroxidase (GPX4), sperm motility (i.e., ADP/ATP translocase 4, ROPN1L, and tektin-5), capacitation (i.e., ROPN1L), and fecundation (i.e., sperm surface protein Sp17 (SPA17)), or were important defense systems against oxidative stress (i.e., nucleoside diphosphate kinase homolog 5 and catalase).

*In silico* protein network analysis indicated that the proteins that were upregulated during the dry (breeding) season, such as ROPN1L, phospholipid hydroperoxide glutathione, HSPA2, and SPA17, interacted with 10 other proteins. Among these interactions, only ROPN1L and phospholipid hydroperoxide glutathione interacted with each other (Fig 2).

**Fig 2.**
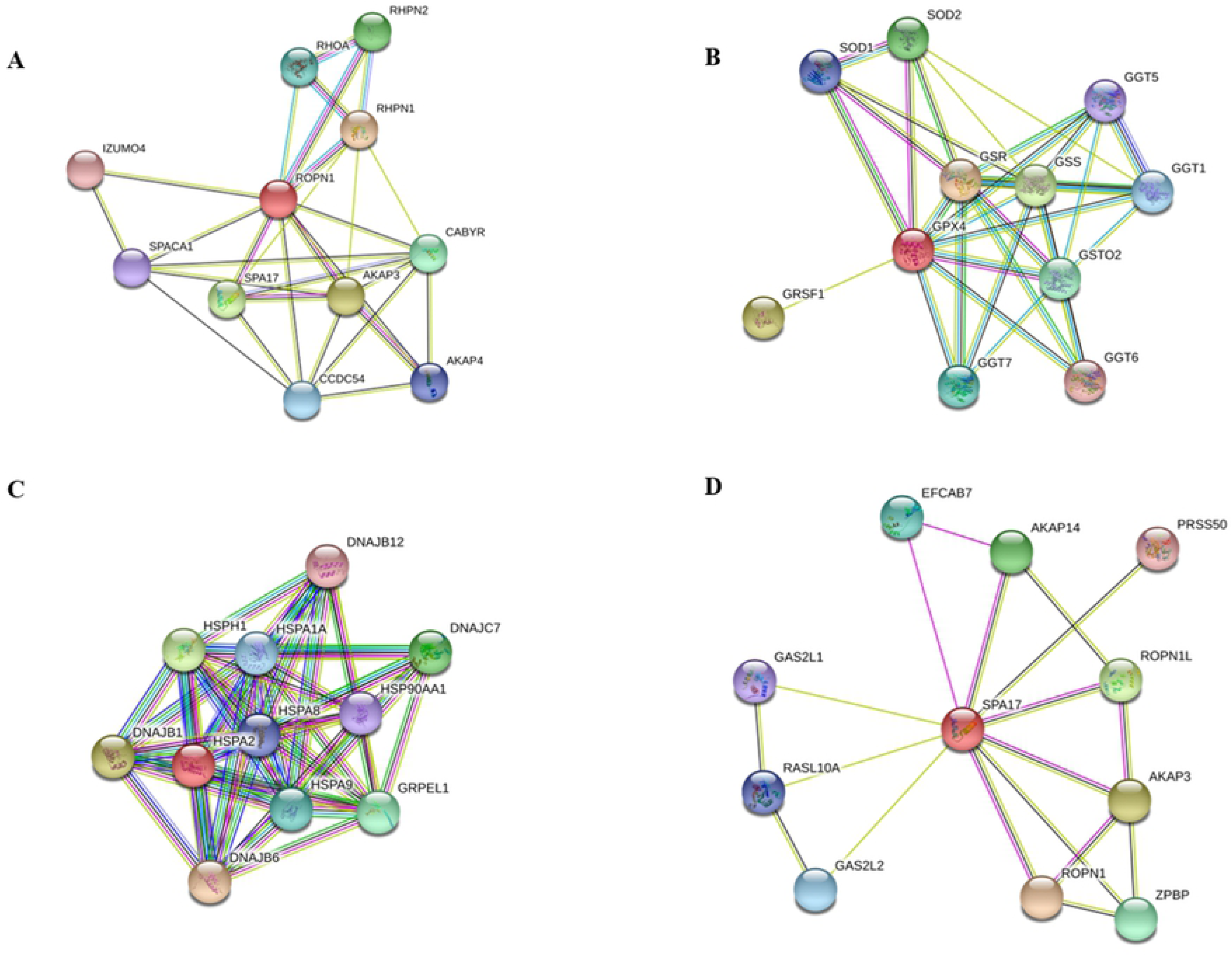
Protein interaction analysis. Proteins were analyzed with the wed-based STRING software. Analyzed proteins were: a-Ropporin-1-like; b-Phospholipid hydroperoxide glutathione; c-Heat shock proteins 70kDa protein 2; d-Sperm surface protein Sp17. Different line color represents the types of evidence for the association. Green textming; black coexpression; blue databases; and pink experiments. AKAP3 A-kinase anchor protein 3; SPA17 Sperm surface protein Sp17; CABYR Calcium-binding tyrosine phosphorylation-regulated protein; RHPN1 Rhophilin-1; RSPH3 Radial spoke head protein 3 homolog; CCDC63 Coiled-coil domain-containing protein 63; TRPV6 Transient receptor potential cation channel subfamily V member 6; DNALI1 Axonemal dynein light intermediate polypeptide 1; TEKT3 Tektin-3; CFAP36 Cilia- and flagella-associated protein 36; GSR Glutathione reductase, mitochondrial; GRSF1 G-rich sequence factor 1; GSS Glutathione synthetase; SOD2 Superoxide dismutase [Mn], mitochondrial; GSTO2 Glutathione S-transferase omega-2; GGT7 Glutathione hydrolase 7; GGT1 Glutathione hydrolase 1 proenzyme; SOD1 Superoxide dismutase [Cu-Zn]; GGT5 Glutathione hydrolase 5 proenzyme; GGT6 Glutathione hydrolase 6; DNAJB6 DnaJ (Hsp40) homolog, subfamily B, member 6; DNAJB1 DnaJ (Hsp40) homolog, subfamily B, member 1; HSPH1 Heat shock 105kDa/110kDa protein 1; DNAJC7 DnaJ (Hsp40) homolog, subfamily C, member 7; GRPEL1 GrpE-like 1, mitochondrial (E. coli); HSPA9 Heat shock 70kDa protein 9 (mortalin); HSPA1A Heat shock 70kDa protein 1A; HSPA8 Heat shock 70kDa protein 8; HSP90AA1 Heat shock protein 90kDa alpha (cytosolic), class A member 1; DNAJB12 DnaJ (Hsp40) homolog, subfamily B, member 12 (409 aa); ROPN1 Rhophilin associated tail protein 1 (212 aa); AKAP3 A kinase (PRKA) anchor protein 3; ROPN1L Rhophilin associated tail protein 1-like (230 aa); AKAP14 A kinase (PRKA) anchor protein 14; ZPBP Zona pellucida binding protein; EFCAB7 EF-hand calcium binding domain 7 (629 aa); GAS2L2 Growth arrest-specic 2 like 2; RASL10A RAS-like, family 10, member A; GAS2L1 Growth arrest-specic 2 like 1; PRSS50 Protease, serine, 50.

**Table 2.**
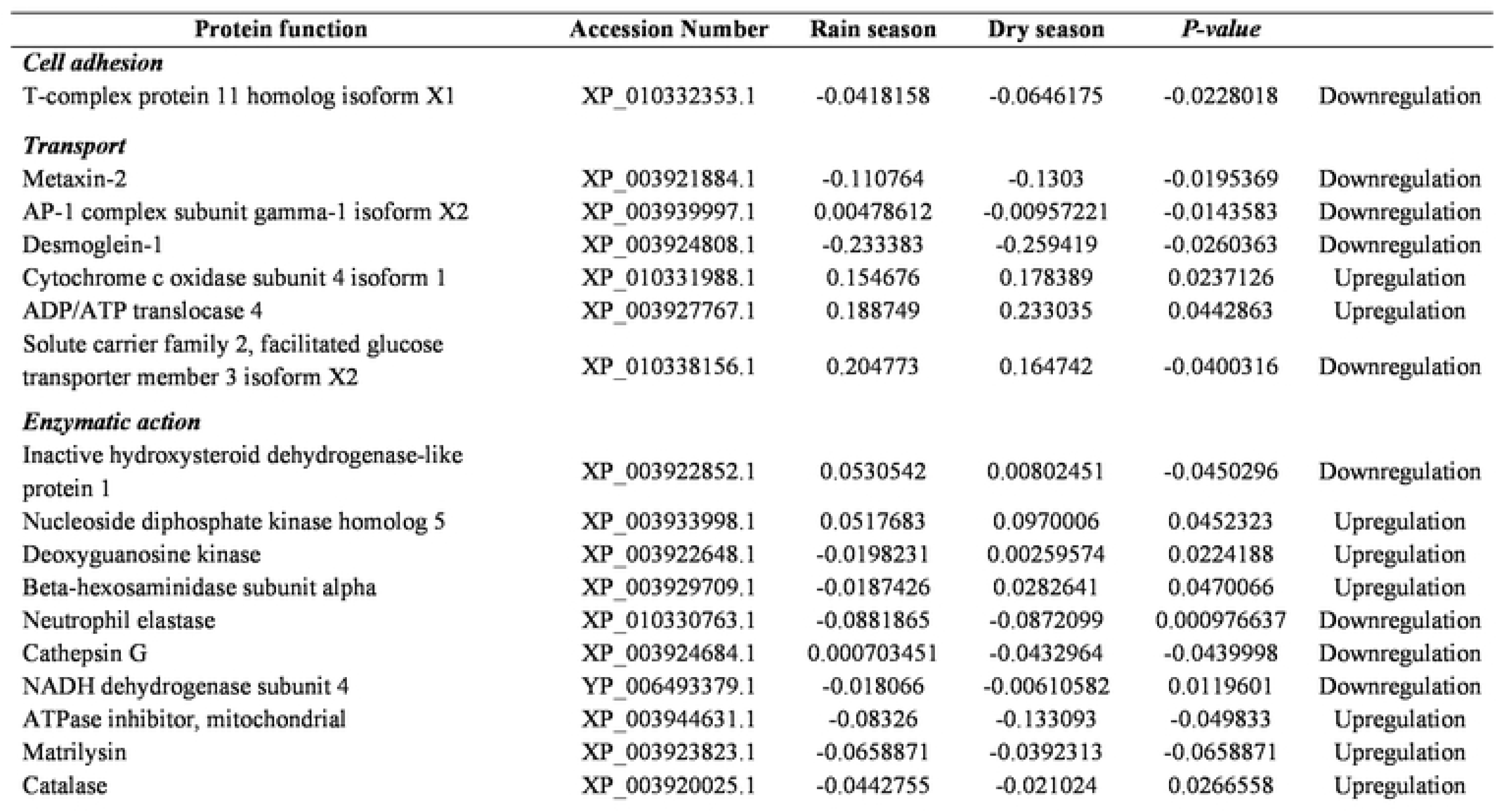
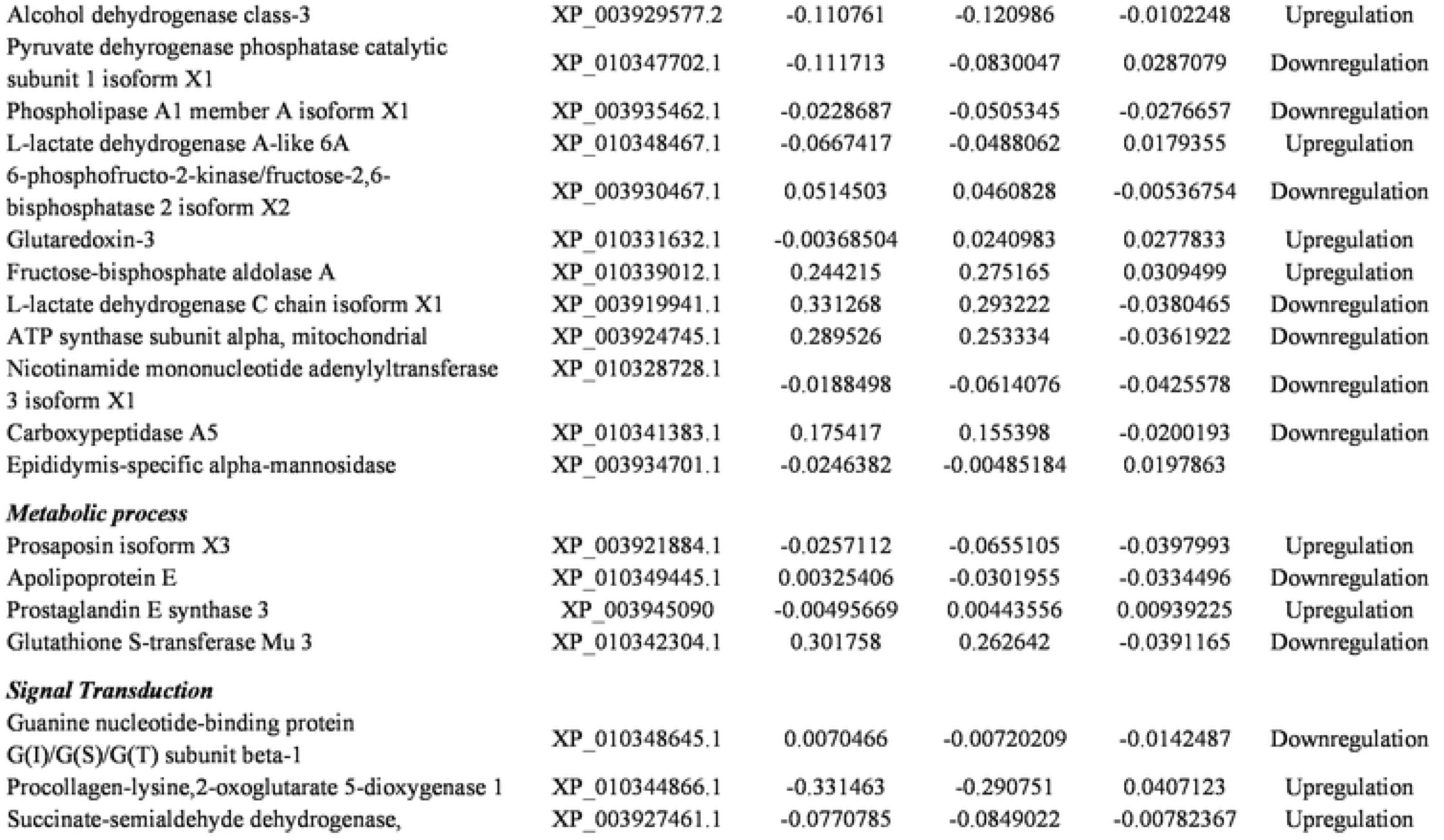
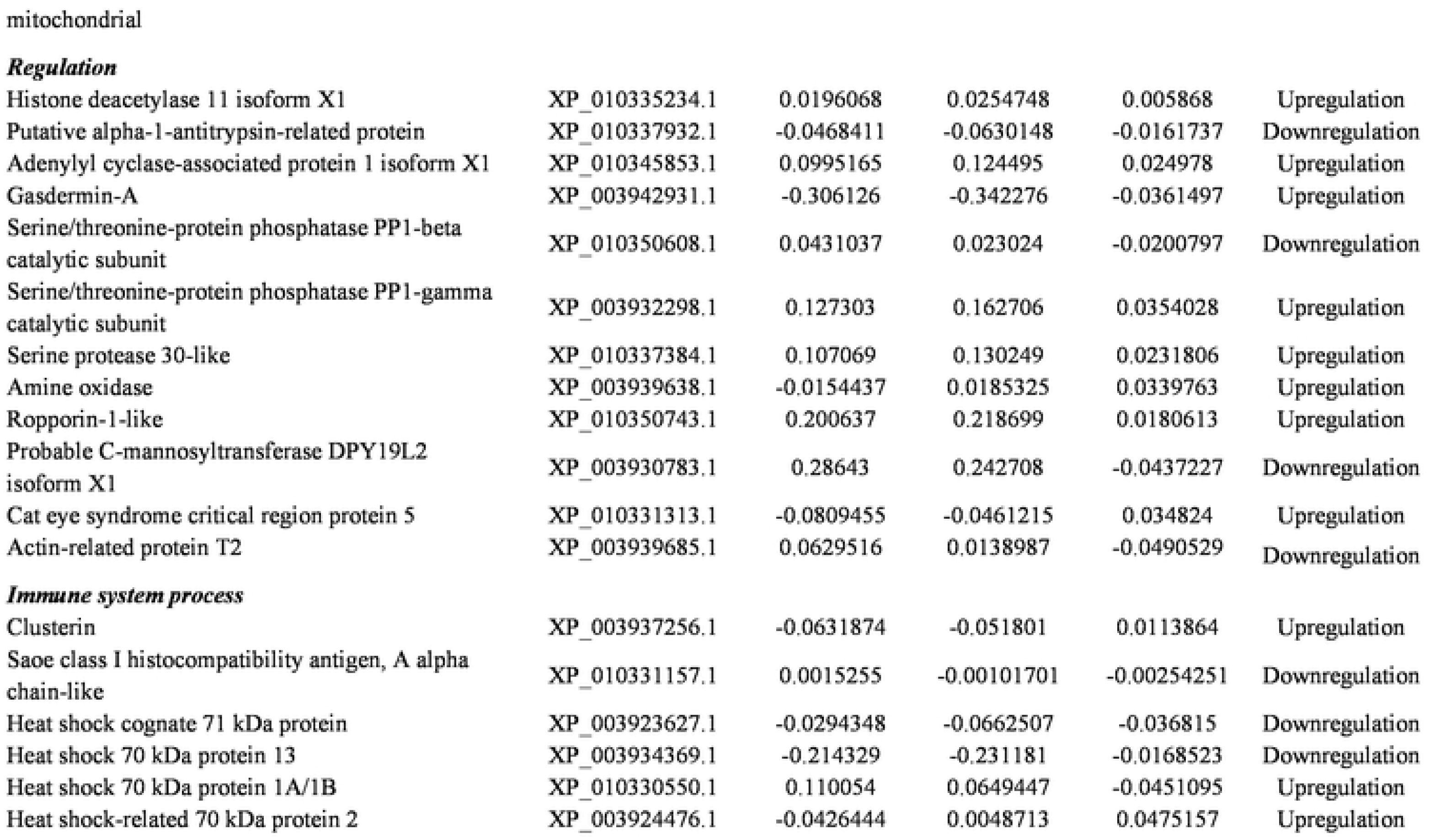
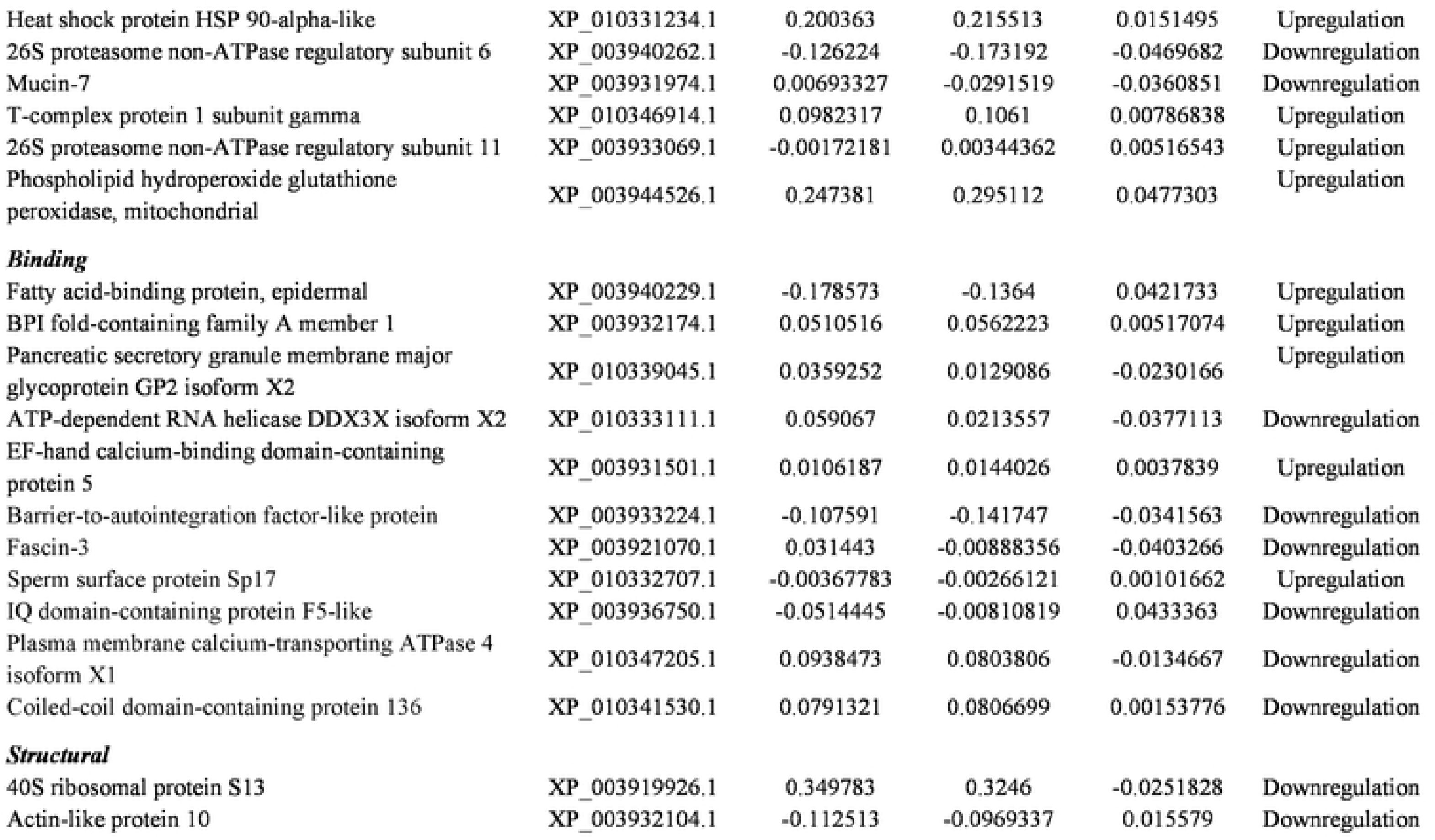
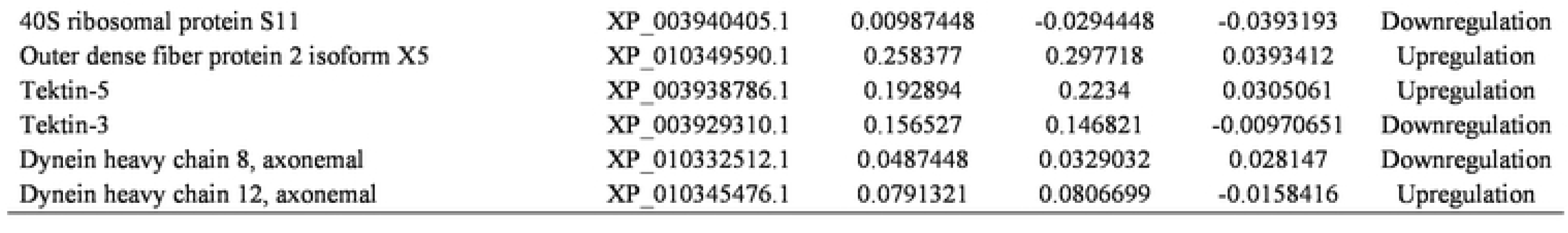
Upregulation or downregulation of sperm protein expression (µg/mL) in *Saimiri collinsi* during the rainy (non-breeding) and dry (breeding) seasons.

| Protein function | Accession Number | Rain season | Dry season | *P-value* |  |
| --- | --- | --- | --- | --- | --- |
| ***Cell adhesion*** |  |  |  |  |  |
| T-complex protein 11 homolog isoform X1 | XP_010332353.1 | -0.0418158 | -0.0646175 | -0.0228018 | Downregulation |
| ***Transport*** |  |  |  |  |  |
| Metaxin-2 | XP_003921884.1 | -0.110764 | -0.1303 | -0.0195369 | Downregulation |
| AP-1 complex subunit gamma-1 isoform X2 | XP_003939997.1 | 0.00478612 | -0.00957221 | -0.0143583 | Downregulation |
| Desmoglein-1 | XP_003924808.1 | -0.233383 | -0.259419 | -0.0260363 | Downregulation |
| Cytochrome c oxidase subunit 4 isoform 1 | XP_010331988.1 | 0.154676 | 0.178389 | 0.0237126 | Upregulation |
| ADP/ATP translocase 4 | XP_003927767.1 | 0.188749 | 0.233035 | 0.0442863 | Upregulation |
| Solute carrier family 2, facilitated glucose transporter member 3 isoform X2 | XP_010338156.1 | 0.204773 | 0.164742 | -0.0400316 | Downregulation |
| ***Enzymatic action*** |  |  |  |  |  |
| Inactive hydroxysteroid dehydrogenase-like protein 1 | XP_003922852.1 | 0.0530542 | 0.00802451 | -0.0450296 | Downregulation |
| Nucleoside diphosphate kinase homolog 5 | XP_003933998.1 | 0.0517683 | 0.0970006 | 0.0452323 | Upregulation |
| Deoxyguanosine kinase | XP_003922648.1 | -0.0198231 | 0.00259574 | 0.0224188 | Upregulation |
| Beta-hexosaminidase subunit alpha | XP_003929709.1 | -0.0187426 | 0.0282641 | 0.0470066 | Upregulation |
| Neutrophil elastase | XP_010330763.1 | -0.0881865 | -0.0872099 | 0.000976637 | Downregulation |
| Cathepsin G | XP_003924684.1 | 0.000703451 | -0.0432964 | -0.0439998 | Downregulation |
| NADH dehydrogenase subunit 4 | YP_006493379.1 | -0.018066 | -0.00610582 | 0.0119601 | Downregulation |
| ATPase inhibitor, mitochondrial | XP_003944631.1 | -0.08326 | -0.133093 | -0.049833 | Upregulation |
| Matrilysin | XP_003923823.1 | -0.0658871 | -0.0392313 | -0.0658871 | Upregulation |
| Catalase | XP_003920025.1 | -0.0442755 | -0.021024 | 0.0266558 | Upregulation |
| Alcohol dehydrogenase class-3 | XP_003929577.2 | -0.110761 | -0.120986 | -0.0102248 | Upregulation |
| Pyruvate dehydrogenase phosphatase catalytic subunit 1 isoform X1 | XP_010347702.1 | -0.111713 | -0.0830047 | 0.0287079 | Downregulation |
| Phospholipase A1 member A isoform X1 | XP_003935462.1 | -0.0228687 | -0.0505345 | -0.0276657 | Downregulation |
| L-lactate dehydrogenase A-like 6A | XP_010348467.1 | -0.0667417 | -0.0488062 | 0.0179355 | Upregulation |
| 6-phosphofructo-2-kinase/fructose-2,6-bisphosphatase 2 isoform X2 | XP_003930467.1 | 0.0514503 | 0.0460828 | -0.00536754 | Downregulation |
| Glutaredoxin-3 | XP_010331632.1 | -0.00368504 | 0.0240983 | 0.0277833 | Upregulation |
| Fructose-bisphosphate aldolase A | XP_010339012.1 | 0.244215 | 0.275165 | 0.0309499 | Upregulation |
| L-lactate dehydrogenase C chain isoform X1 | XP_003919941.1 | 0.331268 | 0.293222 | -0.0380465 | Downregulation |
| ATP synthase subunit alpha, mitochondrial | XP_003924745.1 | 0.289526 | 0.253334 | -0.0361922 | Downregulation |
| Nicotinamide mononucleotide adenylyltransferase 3 isoform X1 | XP_010328728.1 | -0.0188498 | -0.0614076 | -0.0425578 | Downregulation |
| Carboxypeptidase A5 | XP_010341383.1 | 0.175417 | 0.155398 | -0.0200193 | Downregulation |
| Epididymis-specific alpha-mannosidase | XP_003934701.1 | -0.0246382 | -0.00485184 | 0.0197863 |  |
| ***Metabolic process*** |  |  |  |  |  |
| Prosaposin isoform X3 | XP_003921884.1 | -0.0257112 | -0.0655105 | -0.0397993 | Upregulation |
| Apolipoprotein E | XP_010349445.1 | 0.00325406 | -0.0301955 | -0.0334496 | Downregulation |
| Prostaglandin E synthase 3 | XP_003945090 | -0.00495669 | 0.00443556 | 0.00939225 | Upregulation |
| Glutathione S-transferase Mu 3 | XP_010342304.1 | 0.301758 | 0.262642 | -0.0391165 | Downregulation |
| ***Signal Transduction*** |  |  |  |  |  |
| Guanine nucleotide-binding protein G(I)/G(S)/G(T) subunit beta-1 | XP_010348645.1 | 0.0070466 | -0.00720209 | -0.0142487 | Downregulation |
| Procollagen-lysine,2-oxoglutarate 5-dioxygenase 1 | XP_010344866.1 | -0.331463 | -0.290751 | 0.0407123 | Upregulation |
| Succinate-semialdehyde dehydrogenase, | XP_003927461.1 | -0.0770785 | -0.0849022 | -0.00782367 | Upregulation |
| **Regulation** |  |  |  |  |  |
| Histone deacetylase 11 isoform X1 | XP_010335234.1 | 0.0196068 | 0.0254748 | 0.005868 | Upregulation |
| Putative alpha-1-antitrypsin-related protein | XP_010337932.1 | -0.0468411 | -0.0630148 | -0.0161737 | Downregulation |
| Adenylyl cyclase-associated protein 1 isoform X1 | XP_010345853.1 | 0.0995165 | 0.124495 | 0.024978 | Upregulation |
| Gasdermin-A | XP_003942931.1 | -0.306126 | -0.342276 | -0.0361497 | Upregulation |
| Serine/threonine-protein phosphatase PP1-beta catalytic subunit | XP_010350608.1 | 0.0431037 | 0.023024 | -0.0200797 | Downregulation |
| Serine/threonine-protein phosphatase PP1-gamma catalytic subunit | XP_003932298.1 | 0.127303 | 0.162706 | 0.0354028 | Upregulation |
| Serine protease 30-like | XP_010337384.1 | 0.107069 | 0.130249 | 0.0231806 | Upregulation |
| Amine oxidase | XP_003939638.1 | -0.0154437 | 0.0185325 | 0.0339763 | Upregulation |
| Ropporin-1-like | XP_010350743.1 | 0.200637 | 0.218699 | 0.0180613 | Upregulation |
| Probable C-mannosyltransferase DPY19L2 isoform X1 | XP_003930783.1 | 0.28643 | 0.242708 | -0.0437227 | Downregulation |
| Cat eye syndrome critical region protein 5 | XP_010331313.1 | -0.0809455 | -0.0461215 | 0.034824 | Upregulation |
| Actin-related protein T2 | XP_003939685.1 | 0.0629516 | 0.0138987 | -0.0490529 | Downregulation |
| **Immune system process** |  |  |  |  |  |
| Clusterin | XP_003937256.1 | -0.0631874 | -0.051801 | 0.0113864 | Upregulation |
| Saoe class I histocompatibility antigen, A alpha chain-like | XP_010331157.1 | 0.0015255 | -0.00101701 | -0.00254251 | Downregulation |
| Heat shock cognate 71 kDa protein | XP_003923627.1 | -0.0294348 | -0.0662507 | -0.036815 | Downregulation |
| Heat shock 70 kDa protein 13 | XP_003934369.1 | -0.214329 | -0.231181 | -0.0168523 | Downregulation |
| Heat shock 70 kDa protein 1A/1B | XP_010330550.1 | 0.110054 | 0.0649447 | -0.0451095 | Upregulation |
| Heat shock-related 70 kDa protein 2 | XP_003924476.1 | -0.0426444 | 0.0048713 | 0.0475157 | Upregulation |
| Heat shock protein HSP 90-alpha-like | XP_010331234.1 | 0.200363 | 0.215513 | 0.0151495 | Upregulation |
| 26S proteasome non-ATPase regulatory subunit 6 | XP_003940262.1 | -0.126224 | -0.173192 | -0.0469682 | Downregulation |
| Mucin-7 | XP_003931974.1 | 0.00693327 | -0.0291519 | -0.0360851 | Downregulation |
| T-complex protein 1 subunit gamma | XP_010346914.1 | 0.0982317 | 0.1061 | 0.00786838 | Upregulation |
| 26S proteasome non-ATPase regulatory subunit 11 | XP_003933069.1 | -0.00172181 | 0.00344362 | 0.00516543 | Upregulation |
| Phospholipid hydroperoxide glutathione peroxidase, mitochondrial | XP_003944526.1 | 0.247381 | 0.295112 | 0.0477303 | Upregulation |
| ***Binding*** |  |  |  |  |  |
| Fatty acid-binding protein, epidermal | XP_003940229.1 | -0.178573 | -0.1364 | 0.0421733 | Upregulation |
| BPI fold-containing family A member 1 | XP_003932174.1 | 0.0510516 | 0.0562223 | 0.00517074 | Upregulation |
| Pancreatic secretory granule membrane major glycoprotein GP2 isoform X2 | XP_010339045.1 | 0.0359252 | 0.0129086 | -0.0230166 | Upregulation |
| ATP-dependent RNA helicase DDX3X isoform X2 | XP_010333111.1 | 0.059067 | 0.0213557 | -0.0377113 | Downregulation |
| EF-hand calcium-binding domain-containing protein 5 | XP_003931501.1 | 0.0106187 | 0.0144026 | 0.0037839 | Upregulation |
| Barrier-to-autointegration factor-like protein | XP_003933224.1 | -0.107591 | -0.141747 | -0.0341563 | Downregulation |
| Fascin-3 | XP_003921070.1 | 0.031443 | -0.00888356 | -0.0403266 | Downregulation |
| Sperm surface protein Sp17 | XP_010332707.1 | -0.00367783 | -0.00266121 | 0.00101662 | Upregulation |
| IQ domain-containing protein F5-like | XP_003936750.1 | -0.0514445 | -0.00810819 | 0.0433363 | Downregulation |
| Plasma membrane calcium-transporting ATPase 4 isoform X1 | XP_010347205.1 | 0.0938473 | 0.0803806 | -0.0134667 | Downregulation |
| Coiled-coil domain-containing protein 136 | XP_010341530.1 | 0.0791321 | 0.0806699 | 0.00153776 | Downregulation |
| ***Structural*** |  |  |  |  |  |
| 40S ribosomal protein S13 | XP_003919926.1 | 0.349783 | 0.3246 | -0.0251828 | Downregulation |
| Actin-like protein 10 | XP_003932104.1 | -0.112513 | -0.0969337 | 0.015579 | Downregulation |
| 40S ribosomal protein S11 | XP_003940405.1 | 0.00987448 | -0.0294448 | -0.0393193 | Downregulation |
| Outer dense fiber protein 2 isoform X5 | XP_010349590.1 | 0.258377 | 0.297718 | 0.0393412 | Upregulation |
| Tektin-5 | XP_003938786.1 | 0.192894 | 0.2234 | 0.0305061 | Upregulation |
| Tektin-3 | XP_003929310.1 | 0.156527 | 0.146821 | -0.00970651 | Downregulation |
| Dynein heavy chain 8, axonemal | XP_010332512.1 | 0.0487448 | 0.0329032 | 0.028147 | Downregulation |
| Dynein heavy chain 12, axonemal | XP_010345476.1 | 0.0791321 | 0.0806699 | -0.0158416 | Upregulation |

## Discussion

### Proteins associated with spermatogenesis and sperm motility

In *S. collinsi*, it was possible to verify the upregulation of important proteins that participated in spermatogenesis and sperm motility in the dry season (breeding season), such as ROPN1L, HSPA2, cat eye syndrome critical region protein 5, and phospholipid hydroperoxide glutathione. In mice, the loss of ROPN1L impairs sperm motility, cAMP-dependent protein kinase phosphorylation, and fibrous sheath integrity [37]. ROPN1L is a sperm flagellar protein that binds A-kinase anchoring protein (AKAP) 3 and 4, which are primary components of the sperm fibrous sheath. The fibrous sheath is a flagellar cytoskeletal structure unique to sperm that surrounds the outer dense fibers and axoneme [37, 38]. The degradation of AKAP3 and subsequent dephosphorylation of tyrosine result in sperm capacitation [39].

Heat shock proteins (HSPs) are chaperone proteins that are expressed in response to cell stress [40, 41]. Several HSP family members are expressed in the sperm, such as HSP 70 kDa (HSP70), which appears in the acrosome membranes. HSP 60 kDa (HSP60) is located primarily in the sperm midpiece, in association with the mitochondria, whereas HSP 90-alpha (HSP90AA1) is located in the sperm flagellum [42]. HSP60, HSP70, and HSP90AA1 are known components of sperm in different species, such as humans [43], rams [44], bulls, stallions, cats, and dogs [45]. The acrosomal HSP70 has a role in gamete interaction and fertilization [46], whereas HSP90AA1 expression has been correlated with the resistance of sperm to freezing [47, 48] since this protein is characterized as a ubiquitous molecular chaperone that provides protection and protein folding during thermal stress and resistance against cell oxidative stress [49].

HSPA2, which is a molecular chaperone that assists in the folding, transport, and assembly of proteins in the cytoplasm, mitochondria, and endoplasmic reticulum and is a testis-specific member of the 70-kDa family [50], has been suggested to be crucially involved in spermatogenesis and meiosis [51]. In humans, the downregulation of HSPA2 mRNA was observed in testes with abnormal spermatogenesis, and the protein expression was high in normal spermatogenesis and low in spermatogenesis arrest [52]. Human HSPA2 was shown to regulate the expression of the sperm surface receptors involved in sperm-oocyte recognition [53], and its depression in the testes was also associated with spermatogenic impairment and the fertilization rate in men with azoospermia who were treated with intracytoplasmic sperm injections [54].

Sperm motility is driven mainly by the energy produced by the mitochondria present in the intermediate piece of the male gamete [55]. However, the axoneme is another important cellular component that is directly associated with sperm motility. The dynein heavy chains have been annotated as subunits of the axonemal dynein complexes, which are multisubunit axonemal ATPase complexes that generate the force for cilia motility and govern the beat frequency [56]. DNAH1 is related to spermatogenesis and cell proliferation [57]. In humans, mutations in DNAH1 cause multiple morphologic abnormalities of the sperm flagella, leading to male infertility [21]. The radial spoke proteins play a key role in regulating dynein activity and flagellar motility [58, 59].

In this context, Imai et al. [60] showed that the failure to express phospholipid hydroperoxide glutathione peroxidase (GPX4) caused human male infertility, with 30% of men diagnosed with oligoasthenozoospermia showing a significant decrease in the level of the enzyme. Those authors also found a significantly lower number of spermatozoa in the semen and significantly lower motility of the spermatozoa than those seen in fertile men. GPX4 is an intracellular antioxidant that directly reduces peroxidized phospholipids and is strongly expressed in the mitochondria of the testis and spermatozoa. In bulls, GPX4 is considered a unique marker for seminal quality analysis owing to the direct correlation between the selenoperoxidase and the progressive motility of the sperm [61].

### Capacitation and the acrosome reaction

The acrosome, which is a membrane-bound exocytotic vesicle that is located over the anterior portion of the nucleus, contains the hydrolytic enzymes that are required for the acrosome reaction, binding of the zona pellucida (ZP), penetration through the ZP, and sperm–egg membrane fusion, all of which are indispensable events during the fertilization process [62]. In the acrosome membrane (internal and external membranes), the sperm acrosome membrane-associated family (i.e., SPACA3, SPACA1, and SPACA4) [63, 64] are sperm surface membrane proteins that may be involved in the adhesion and fusion of the sperm to the egg prior to fertilization [65]. SPACA1 and SPACA3 are localized in the acrosomal matrix, including the principal segment and equatorial segment, and are proteins characterized as membrane antigens [63, 65, 66]. SPACA1 may be involved in sperm fusion with the oölemma, since treatment of human sperm with the anti-SPACA1 antibody prevented sperm penetration into zona-free hamster eggs [63]. Fujihara et al. [67] demonstrated that the SPACA1 protein was indispensable for the normal shaping of the sperm heads during spermiogenesis in mice. In humans, this protein was identified as a sperm membrane antigen, with a molecular mass ranging from 32 to 34 kDa [63].

### Sperm–egg fusion

Membrane fusion is a key event in the fertilization process that culminates in the merger of the male–female gamete membranes and cytoplasm and fusion of the genomes, thereby initiating embryonic development [68]. In humans, a change in the expression of the sperm proteins may be a major cause of subfertility in men with normozoospermia [69]. In this context, research has been focusing on the identification of the key molecular players and their functions, and several proteins in the egg or the spermatozoa have been found to be essential for fertilization.

Until now, IZUMO1 has been found to be the essential protein on the sperm side for the fusion process. As a testis-specific protein, IZUMO was discovered on the equatorial segment of the acrosome-reacted mouse spermatozoa through proteomic analysis of the antigen recognized by the monoclonal anti-mouse sperm antibody [70]. IZUMO is present in both mouse (∼56 kDa protein) and human (∼38 kDa protein) sperm [71]. In mice, immunization with the IZUMO protein caused a contraceptive effect in females, which was due to the significantly inhibited fusion of sperm to the zona-free mouse eggs with the anti-PrimeB antibody. However, no effect on sperm motility was observed [72]. IZUMO2, IZUMO3, and IZUMO4 have significant homology with the N-terminal domain of IZUMO1 [73]. Inoue et al. [24] showed the interaction between angiotensin-converting enzyme-3 located on the sperm acrosomal cap and IZUMO1 in the fertilization process. However, it was reported that angiotensin-converting enzyme-3 disappears from the membrane after the acrosome reaction. Nevertheless, the *in silico* protein interaction analysis of IZUMO1 revealed its association with the CD9 molecule, folate receptor 4 (delta) homolog (mouse), folate receptor 1 (adult), folate receptor 2 (fetal), SPACA1, SPACA4, IZUMO family member 4, zona pellucida binding protein 2, and metallopeptidase domain 2.

After the acrosome reaction, the C-terminal calmodulin domain (20 kDa) of SPA17 (located on the external side of the sperm plasma membrane) is proteolytically cleaved to 17 kDa and then binds to the extracellular matrix of the oocyte. This C-terminus of SPA17 plays a role in cell–cell adhesion [74, 75]. In our study, SPA17 was shown to be upregulated during the dry season, implying that this protein could also be involved in the fertilization processes in the breeding season of *S. collinsi*.

Our results on the seminal quality also showed that proteomics is an important complementary tool for use toward understanding and elucidating the influence of seasonality on the sperm cells in *S. collinsi*, since it was not possible to verify this influence by classic seminal analysis for this species. Additionally, it is important to mention that the results of the seminal parameters analyzed (viz., appearance, semen volume, pH, and sperm concentration, motility, vigor, and morphology) were similar to those previously reported for fresh Amazon squirrel monkey sperm (liquid fraction) and sperm from the coagulated fraction after dilution in ACP-118 [2–4]. However, this was the first time that a comparison of these parameters during the dry and rainy seasons was performed for this species.

Although the seminal pH was higher in the dry season, it was slightly alkaline during both seasons and similar to the range reported elsewhere for *S. collinsi* (pH 6.5–8.0) [2–4] as well as for the Neotropical primates *Alouatta caraya* (pH 8.1) [76], *Ateles geoffroyi* (pH 8.0) [77], *Callithrix jacchus* (pH 7.4–7.6) [78, 79], *Callithrix penicillata* [80], and *Callimico goeldii* (pH 6.1) [81]. In women, the acidic vaginal environment is toxic to sperm because the optimal pH for sperm viability ranges from 7.0 to 8.5, and a reduction in sperm motility is seen at a pH of less than 6.0. However, during human sexual intercourse, the vaginal epithelium produces a transudate that lubricates the vagina and elevates the vaginal pH to 7.0 [82]. This physiological modification to accommodate the alkaline pH of semen temporarily protects the spermatozoa and creates an optimal environment in the cervix for sperm motility [83].

It is worth mentioning that measurement of the seminal pH in our study was only possible with the liquid fraction, as it was necessary to dilute the coagulated fraction in order to establish its pH value. The ACP-118 extender used for non-human primates, including species of the genus *Saimiri* [2–4], has a pH (6.5) that is compatible to the liquid fraction of *S. collinsi* semen. ACP-118 is composed of different bioactive enzymes (e.g., phosphatase, catalase, and dehydrogenase), which may support coagulum liquefaction. This extender also contains ascorbic acid and polyphenol oxidases, which are antioxidants that maintain the sperm quality during and after incubation [84, 85]. In this way, the ACP-118 composition may have affected the quality of the *S. collinsi* sperm, since there was no difference in the sperm parameters analyzed between the dry and rainy seasons. Thus, our results showed that the ACP-118 extender used for coagulum liquefaction was able to maintain similar sperm qualities in both seasonal periods.

## Conclusions

The present study is a comprehensive overview of the sperm proteome in the Amazon squirrel monkey, and is the broadest inventory (investigation) of the sperm proteome in the genus *Saimiri* as well as in Neotropical primates thus far. The knowledge acquired about the sperm proteins is a significant step forward in helping toward our understanding of the reproductive biology of the genus *Saimiri*, as it provides crucial information for the elucidation of the underlying mechanisms associated with sperm function. In this way, our study amplifies the advances in biotechnological research on animal reproduction for the conservation of endangered species, and provides a reference for similar studies on other Neotropical primates. Nevertheless, further studies should be carried out to verify the differences in the patterns of protein expression throughout the year in other species of the genus *Saimiri*.

## Acknowledgments

The authors thank Coordenação de Aperfeiçoamento de Pessoal de Nível Superior - Brasil (CAPES) - Finance Code 001, and Conselho Nacional de Desenvolvimento Científico e Tecnológico (Projeto Universal 01-2016/ Processo No. 421649/2016-0) for their financial support. We would also like to thank the National Primate Center (Conselho Executivo das Normas-Padrão, Brazil) and Norwegian University of Life Sciences (Norway) for the technical support provided during this research.

## Supporting information

S1 Table. Spectral count of *Saimiri collinsi* sperm protein throughout an entire year (.XLS).

S2 Table. Sperm proteins of *Saimri collinsi* that participate in important reproductive events (.XLS).

S3 Table. Two-sample tests of the sperm protein concentrations in *Saimiri collinsi* during the dry and rainy seasons.

